# Flotillin-dependent lipid-raft microdomains are required for functional phagolysosomes against fungal infections

**DOI:** 10.1101/606939

**Authors:** Franziska Schmidt, Andreas Thywißen, Marie Röcker, Cristina Cunha, Zoltán Cseresnyés, Hella Schmidt, Silvia Galiani, Markus H. Gräler, Georgios Chamilos, João F. Lacerda, António Campos, Christian Eggeling, Marc Thilo Figge, Thorsten Heinekamp, Scott G. Filler, Agostinho Carvalho, Axel A. Brakhage

**Affiliations:** Department of Molecular and Applied Microbiology, Leibniz Institute for Natural Product Research and Infection Biology – Hans Knöll Institute (HKI), 07745 Jena, Germany; Department of Microbiology and Molecular Biology, Institute of Microbiology, Friedrich Schiller University Jena, 07745 Jena, Germany; Life and Health Sciences Research Institute (ICVS), School of Medicine, University of Minho, Campus de Gualtar, 4710-057 Braga, Portugal; ICVS/3B’s – PT Government Associate Laboratory, Braga/Guimarães, Portugal; Research Group Applied Systems Biology, Leibniz Institute for Natural Product Research and Infection Biology (HKI), 07745 Jena, Germany; MRC Human Immunology Unit, Weatherall Institute of Molecular Medicine, University of Oxford, Headley Way, Oxford OX3 9DS, UK; Department of Anesthesiology and Intensive Care Medicine, Center for Sepsis Control and Care (CSCC), and the Center for Molecular Biomedicine (CMB), University Hospital Jena, 07740 Jena, Germany; Department of Medicine, University of Crete, and Institute of Molecular Biology and Biotechnology, Foundation for Research and Technology, 71300 Heraklion, Crete, Greece; Serviço de Hematologia e Transplantação de Medula, Hospital de Santa Maria, 1649-035 Lisboa, Portugal, and Instituto de Medicina Molecular, Faculdade de Medicina de Lisboa, 1649-028 Lisboa, Portugal; Serviço de Transplantação de Medula Óssea (STMO), Instituto Português de Oncologia do Porto, 4200-072 Porto, Portugal; Institute of Applied Optics, Friedrich Schiller University Jena, and Department of Biophysical Imaging, Leibniz Institute of Photonic Technology (IPHT), Jena, Germany; Division of Infectious Diseases, Los Angeles Biomedical Research Institute at Harbor-UCLA Medical Center, Torrance CA 90502, USA, David Geffen School of Medicine at University of California, Los Angeles, Los Angeles, California, USA

**Keywords:** *Aspergillus fumigatus*, conidial melanin, phagolysosome, lipid rafts, flotillin, vATPase, NADPH oxidase, calcium signaling, macrophage, SNP, fungal pathogenesis

## Abstract

Lipid rafts form signaling platforms on biological membranes with incompletely characterized role in immune response to infection. Here we report that lipid raft microdomains are essential components of the phagolysosomal membrane of macrophages. Genetic deletion of the lipidraft chaperons flotillin-1 and flotillin-2 demonstrate that the assembly of both major defense complexes vATPase and NADPH oxidase on the phagolysosomal membrane requires lipid rafts. Furthermore, we discovered a new virulence mechanism leading to the dysregulation of lipid-raft formation by melanized wild-type conidia of the important human-pathogenic fungus *Aspergillus fumigatus*. This results in reduced phagolysosomal acidification. Phagolysosomes with ingested melanized conidia contain a reduced amount of free Ca^2+^ ions as compared to phagolysosomes with melanin-free conidia. In agreement with a role of Ca^2+^ for generation of functional lipid rafts, we show that Ca^2+^-dependent calmodulin activity is required for lipid-raft formation on the phagolysosome. We identified a single nucleotide polymorphism in the human *FLOT1* gene that results in heightened susceptibility for invasive aspergillosis in hematopoietic stem-cell transplant recipients. Collectively, flotillin-dependent lipid rafts on the phagolysosomal membrane play an essential role in protective antifungal immunity in humans.

## INTRODUCTION

Many bacteria manipulate professional phagocytes to counteract immune responses. For example, *Mycobacterium spp., Legionella spp*., and *Listeria spp*., withstand intracellular degradation after ingestion by phagocytes (Cambier, Falkow et al., 2014, Carvalho, Sousa et al., 2014, Ensminger, 2016). This is accomplished by their ability to interfere with the formation of a functional hostile phagolysosome (PL). Pathogenic fungi have also developed mechanisms to avoid being killed after they are taken up by phagocytes. The encapsulated yeast, *Cryptococcus neoformans*, damages phagosomal membranes, deploys potent antioxidant mechanisms and manipulates phagosomal maturation, which enables the fungus to survive and replicate inside phagosomes (Levitz, Nong et al., 1999, Smith, Dixon et al., 2015, Tucker & Casadevall, 2002, Zaragoza, Chrisman et al., 2008). Furthermore, *C. neoformans* can induce non-lytic exocytosis, in which the ingested pathogen is expelled from the phagocyte without lysing it (Alvarez & Casadevall, 2006, Ma, Croudace et al., 2006, Nicola, Robertson et al., 2011). Other fungi, such as the yeast forms of *Candida spp*. and *Histoplasma capsulatum* employ mechanisms to establish an intracellular lifecycle or to avoid killing inside the PL, respectively (Fernandez-Arenas, Bleck et al., 2009, Newman, Gootee et al., 2006, Seider, Brunke et al., 2011, Strasser, Newman et al., 1999). For example, *H. capsulatum* yeast cells reduce phagosomal acidification by releasing the saposin-like protein CBP that reduces vacuolar ATPase (vATPase) accumulation on the phagosomal membrane (Batanghari, Deepe et al., 1998, Woods, 2003). *Rhizopus spp*. establish intracellular persistence inside alveolar macrophages. Lack of intracellular swelling of *Rhizopus* conidia results in surface retention of melanin, which induces phagosome maturation arrest through inhibition of LC3-associated phagocytosis (LAP) (Andrianaki, Kyrmizi et al., 2018).

In line, we previously reported that dihydroxynaphthalene (DHN) melanin of conidia (spores) of the important pathogenic fungus *Aspergillus fumigatus* that is predicted to cause more than 200.000 life-threatening infections worldwide (Brown, Denning et al., 2012, Kosmidis & Denning, 2015), is responsible for establishing a non-hostile intracellular niche inside phagocytes by inhibiting phagolysosomal acidification, and as a consequence killing of conidia, LAP and apoptosis of phagocytes (Akoumianaki, Kyrmizi et al., 2016, Heinekamp, Thywissen et al., 2012, Schmidt, Vlaic et al., 2018, Thywissen, Heinekamp et al., 2011, Volling, Thywissen et al., 2011). The grey-green surface pigment consisting of dihydroxynaphthalene (DHN) melanin (Jahn, Koch et al., 1997, Langfelder, Streibel et al., 2003, Tsai, Chang et al., 1998) plays a pivotal role in this process because melanin ghosts, *i.e*., isolated pigment particles, led to reduced acidification to the same extent as wild-type conidia and impaired LAP, whereas conidia from the pigmentless *pksP* mutant showed full acidification of PLs and full activation of LAP (Akoumianaki et al., 2016, Thywissen et al., 2011). Furthermore, the absence of conidial melanin in the *pksP* mutant results in a higher phagocytosis rate, faster endocytotic processing, and increased production of pro-inflammatory cytokines and chemokines by phagocytes most likely due to better recognition of *pksP* conidia by the dectin-1 receptor (Chai, Netea et al., 2010, Luther, Torosantucci et al., 2007, Mech, Thywissen et al., 2011, Thywissen et al., 2011). Further data indicated that DHN-melanin is able to abrogate the release of free Ca^2+^ ions to the peri-phagosomal area thereby inhibiting calmodulin activity and impairing phagosome function (Kyrmizi, Ferreira et al., 2018). However, the downstream effector pathways regulating phagosome biogenesis and pathogen killing are incompletely characterized.

For fungal pathogens, the molecular mechanisms involved in the dysregulation of PLs and, furthermore, many aspects of phagosome maturation are unsolved. Here, we add novel insight into both aspects by the finding of lipid-raft microdomains as being essential for phagosome maturation, defense against invasive aspergillosis and also as a target of pathogens. We discovered an unprecedented virulence mechanism depending on the inhibition of flotillin-dependent lipid raft formation in phagolysosomal membranes by conidial melanin.

## RESULTS

### Reduced phagolysosomal acidification increases conidia-induced damage of macrophages

If a macrophage ingests an *A. fumigatus* conidium and fails to kill it, the conidium swells and germinates to produce a hypha that then pierces the macrophage and eventually kills it. We used the vATPase inhibitor, bafilomycin A1, to determine whether inhibition of phagolysosomal acidification affects the extent of macrophage damage. Wild-type conidia caused considerable macrophage damage whereas *pksP* mutant conidia lacking DHN-melanin caused significantly lower host cell damage (Fig. 1A). Abolishing phagolysosomal acidification with bafilomycin A1 increased the macrophage damage caused by both wild-type and pigmentless *pksP* conidia. Importantly, in the presence of bafilomycin A1, wild-type and *pksP* conidia caused virtually identical host cell damage. These results indicate that the capacity of wild-type *A. fumigatus* conidia to reside in a PL with reduced acidification enhances the capacity of the organism to survive within the phagocyte and eventually to kill it.

**Figure 1.**
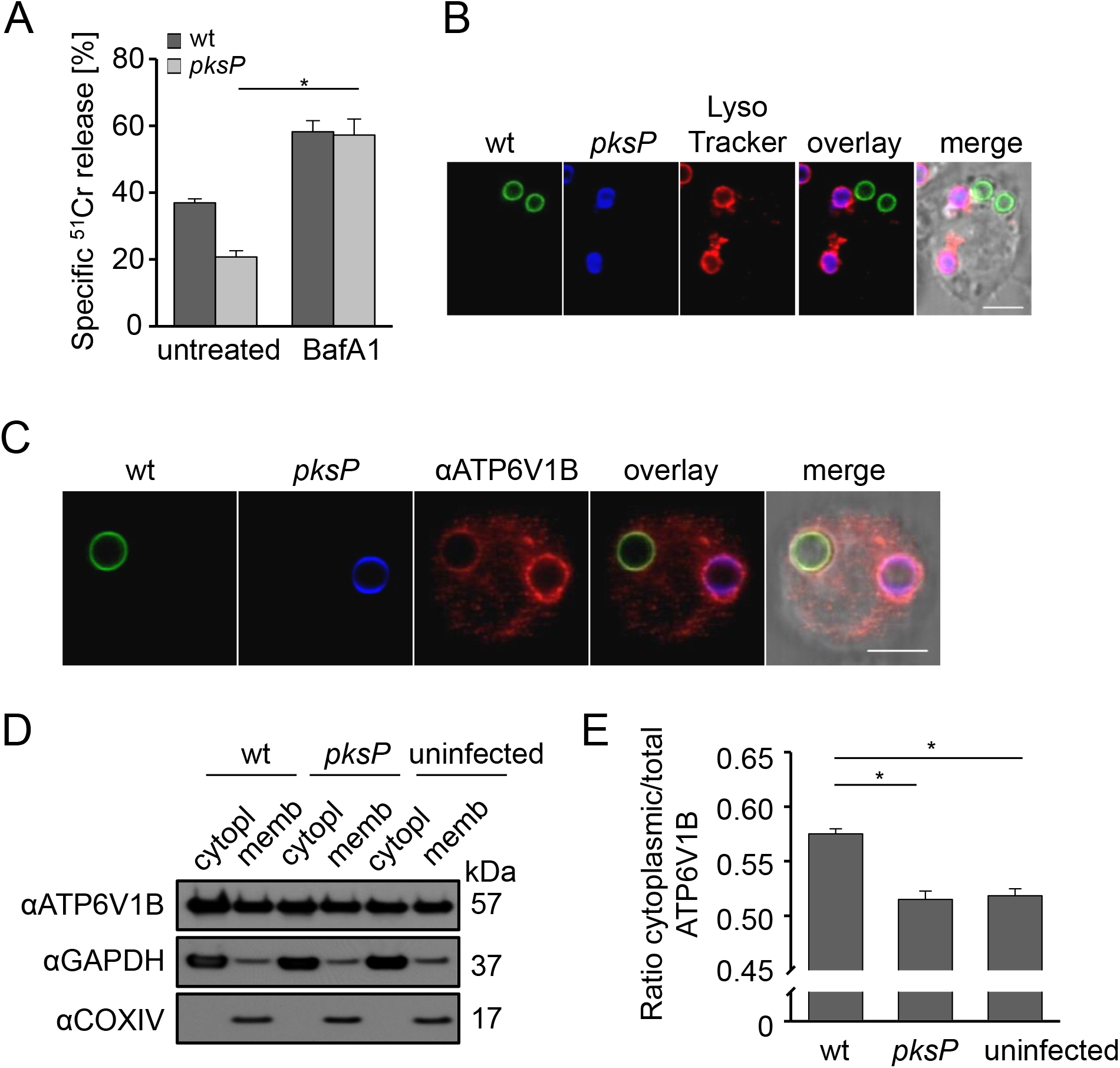
*A. fumigatus* melanized wild-type conidia increase host cell damage and reduce functionality of PLs in which they reside. **A)** Cell damage monitored by ^51^Cr release from RAW264.7 macrophages infected with *A. fumigatus* wild-type or *pksP* conidia; data are represented as mean ± SD; *p* < 0.05. **B)** RAW264.7 macrophage cell had simultaneously phagocytosed melanized wild-type conidia (labeled with FITC, green) and non-pigmented *pksP* conidia (labeled with CFW, blue). Acidified phagolysosome stained with LysoTracker Red. Scale bar = 5 μm **C)** RAW264.7 macrophage had simultaneously phagocytosed a wild-type conidium (FITC-labeled) and a *pksP* conidium (CFW-labeled). Localization of cytoplasmic vATPase subunit V_1_ to the phagolysosomal membrane was monitored by immunofluorescence. Scale bar = 5 μm **D)** Western blot analysis for detection of vATPase V_1_ subunit using an ATP6V1B antibody. Cytoplasmic and membrane fractions of macrophages infected with wild-type or *pksP* conidia were analyzed. As controls, the cytoplasm and membrane specific proteins GAPDH and COXIV, respectively, were detected. **E)** Western blot data (from D) were quantified densitometrically to determine the ratio of cytoplasmic to total ATP6V1B subunit content of RAW264.7 macrophages infected with wildtype or *pksP* conidia; data show mean values ± SD; *p* < 0.05.

### Reduced acidification is specific for PLs containing melanized conidia and correlates with reduced assembly of functional vATPase in the phagolysosomal membrane

To investigate whether the inhibition of phagolysosomal acidification by wild-type conidia is a global phenomenon or is limited to PLs that contain melanized conidia, macrophages were loaded with LysoTracker Red, simultaneously infected with melanized wild-type and pigmentless *pksP* conidia. Acidified PLs were identified by their red fluorescence. We found that when wildtype and *pksP* conidia were ingested by the same macrophage, there was no acidification of the PL containing a wild-type conidium, but clear acidification of the PL containing the *pksP* conidium (Fig. 1B).

We hypothesized that wild-type conidia-containing phagolysosomal membranes contain less active vATPase. Active vATPase consists of two complexes; the V_0_ complex is integrated into the membrane and, when assembled with the cytoplasmic V_1_ complex, shuttles protons across the lipid bilayer (Cotter, Stransky et al., 2015). To analyze the binding of cytoplasmic V_1_ to the V_0_ complex the localization of the V_1_ complex to the membranes of PLs that contained either wildtype or *pksP* conidia was monitored by immunofluorescence. In the same macrophage, we only found faint staining of the phagolysosomal membrane that surrounded wild-type conidia (Fig. 1C). By contrast, there was a strong signal for V_1_ at the phagolysosomal membrane surrounding *pksP* conidia. To further quantify the amount of assembled vATPase, immunoblotting was used to determine the relative amount of the V1 complex in the membrane and cytosolic fractions of macrophages containing wild-type conidia and *pksP* conidia to calculate the ratio of cytosolic V_1_ to the total amount of V_1_ subunits. There was more of the V_1_ complex in the cytosolic fraction in macrophages infected with wild-type conidia relative to macrophages infected with *pksP* conidia (Fig. 1D, E). These results indicate that wild-type conidia-containing PLs contain less assembled vATPase complex and thus reduced acidification. This mechanism is specific to the PL in which melanized conidia are located.

### Melanized conidia interfere with formation of lipid-raft microdomains in PLs

To examine whether the formation of lipid-raft microdomains, which are rich in cholesterol and sphingolipids (Lingwood & Simons, 2010), in the PL membrane was disturbed when DHN-melanin-containing conidia were ingested, the presence of several lipid-raft marker lipids was analyzed. For this purpose, macrophages were infected with melanized and *pksP* conidia and then stained with fluorescently labeled cholera toxin subunit B (CTB) to label the sphingolipid GM1 ganglioside, whose accumulation at distinct membrane sites is characteristic of lipid rafts and which was used as an initial indicator for lipid-raft microdomains (Ledeen & Wu, 2015). Interestingly, in both MH-S and RAW264.7 macrophages that had ingested melanized conidia, there was only minimal CTB staining of the phagolysosomal membrane, despite the strong CTB staining of the macrophage cytoplasmic membrane (Fig. 2A, B). By contrast, in macrophages that had ingested *pksP* conidia, distinct CTB staining of both the phagolysosomal and cytoplasmic membranes was visible (Fig. 2A, B).

**Figure 2.**
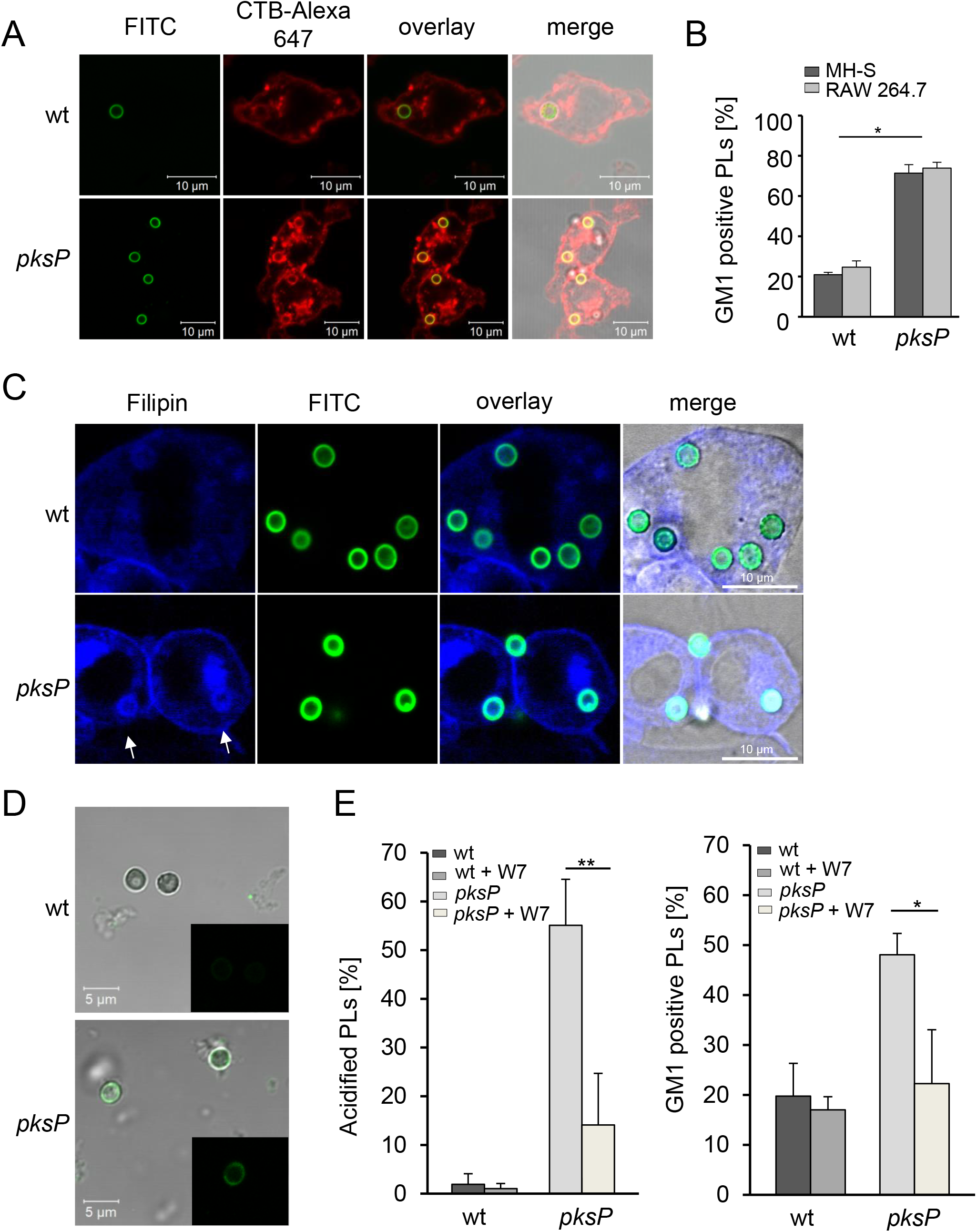
*A. fumigatus* conidia interfere with phagolysosomal lipid-raft formation. **A)** Murine RAW264.7 macrophages stained for GM1 ganglioside (CTB-Alexa 647, red) were infected with FITC-labeled wild-type or *pksP* conidia. **B)** Quantification of GM1 positive phagolysosomes (PLs) in MH-S and RAW264.7 macrophages after infection with wild-type or *pksP* conidia. Data are represented as mean ± SD; *p* < 0.05. **C)** Cholesterol in conidia-containing PLs of RAW264.7 macrophages was detected by filipin III staining. Conidia were stained with FITC; *pksP* conidia-containing PLs display a distinct ring-like structure of cholesterol (indicated by white arrows), which was lacking in wild-type conidia-containing PLs. See also Figures S1, S2, S3 and S4 and supplemental videos 1 and 2. D) Ca^2+^ ions in PLs containing wild-type and *pksP* conidia stained with Fluo-4 AM. **E)** Acidification of PLs and GM1 staining of PLs containing the indicated conidia upon addition of calmodulin inhibitor W7.

To further substantiate this finding, the cholesterol-specific fluorescent probe filipin III and the lipophilic dye DiD were used. As shown by confocal laser scanning microscopy (CLSM), both filipin III (Fig. 2C) and DiD (Fig. S1) weakly labeled membranes surrounding melanized conidia while strongly labeling membranes enclosing *pksP* conidia. These results confirm that the presence of DHN-melanin on the ingested conidium affects the lipid composition of the phagolysosomal membrane.

To determine the relationship between lipid rafts and phagolysosomal acidification, we demonstrated that PLs containing melanized conidia were neither acidified nor contained GM1-stained lipid-raft microdomains, whereas PLs containing *pksP* conidia contained acidified endosomal and lysosomal vesicles that strongly colocalized with lipid-raft microdomains (Fig. S2). The accumulation of lipid-raft microdomains in phagolysosomal membranes enclosing *pksP* conidia was also visualized as 3D reconstruction (supplemental Video S1). By contrast, such lipid-raft spots were less prominent in phagolysosomal membranes enclosing melanized wildtype conidia (supplemental Video S2).

In additional experiments, we found that melanin ghosts had the same capacity as wild-type conidia to reside in non-acidified PLs with reduced formation of lipid-raft microdomains (Fig. S3). Collectively, these data indicate that lipid-raft microdomain formation correlates with the acidification of the PL and that these processes are reduced in PLs containing melanized conidia or melanin ghosts.

To determine whether reduced lipid raft formation influences PL acidification, macrophages were treated with methyl-β-cyclodextrin (MβCD), which depletes the cells of cholesterol and consequently blocks the formation of lipid rafts (Keller & Simons, 1998). MβCD treatment reduced the total cellular cholesterol content to less than 20 % compared to the untreated control (Fig. S4). Exposure to MβCD resulted in a complete absence of acidification of the PL as well as almost all of the endosomal and lysosomal vesicles (Fig. S4). Although approximately 10 % of PLs containing melanized and *pksP* conidia still stained positive for GM1, acidification of these PLs was almost completely abolished (Fig. S4). Thus, the presence of lipid-raft microdomains in the phagolysosomal membrane contributes to proper acidification.

### Inhibition of calmodulin activity reduces microdomain formation in the phagolysosomal membrane

Previously we showed that melanized conidia reduce the release of free Ca^2+^ ions from the phagosome lumen to the peri-phagosomal area, thereby preventing activation of calmodulin and LAP (Kyrmizi et al., 2018). Thus, it was conceivable that Ca^2+^ sequestering by DHN-melanin also dysregulates lipid-raft microdomain formation. To test this assumption we first tested the presence of Ca^2+^ ions in PLs containing melanized and *pksP* conidia using Fluo-4 AM. Fluorescence microscopy indicated a much higher content of Ca^2+^ ions in *pksP*-containing PLs compared with wild-type conidia-containing PLs (Fig. 2D). Next, we inhibited calmodulin activity by addition of the inhibitor W7 that led to reduced formation of lipid-raft microdomains and reduced acidification of pksP-containing PLs (Fig. 2E). Our data suggest that the activation of calmodulin contributes to lipid-raft microdomain formation and its inhibition by the capability of DHN-melanin to sequester Ca^2+^ reduced microdomain formation.

### Lipid-raft microdomains of RAW264.7 macrophages contain flotillin but not caveolin

The role of lipid rafts for phagosome biogenesis and their composition in the phagosomal membrane are unknown. Nevertheless, the differentiation of membrane microdomains into subclasses of lipid-raft domains and caveolae domains is widely accepted (Lingwood & Simons, 2010, Pike, 2009). In RAW264.7 macrophages, we did not detect caveolin-1 or caveolin-2 by immunofluorescence analysis or immunoblotting studies of whole cell extracts (Fig. 3A). Both caveolin-1 and caveolin-2 were detected in control HeLa cells (Fig. 3A). By contrast, flotillin-1, a marker for a different subset of lipid rafts and assumed to act as scaffolding protein that stabilizes lipid-raft microdomains (Banning, Tomasovic et al., 2011, Otto & Nichols, 2011, Stuermer, 2011), colocalized with GM1 ganglioside on phagolysosomal membranes of RAW264.7 macrophages containing *pksP* conidia but not melanized conidia (Fig. 3B, C). These data indicate that *pksP* but not wild-type conidia reside in PLs containing flotillin-dependent lipidraft microdomains in the membrane.

**Figure 3.**
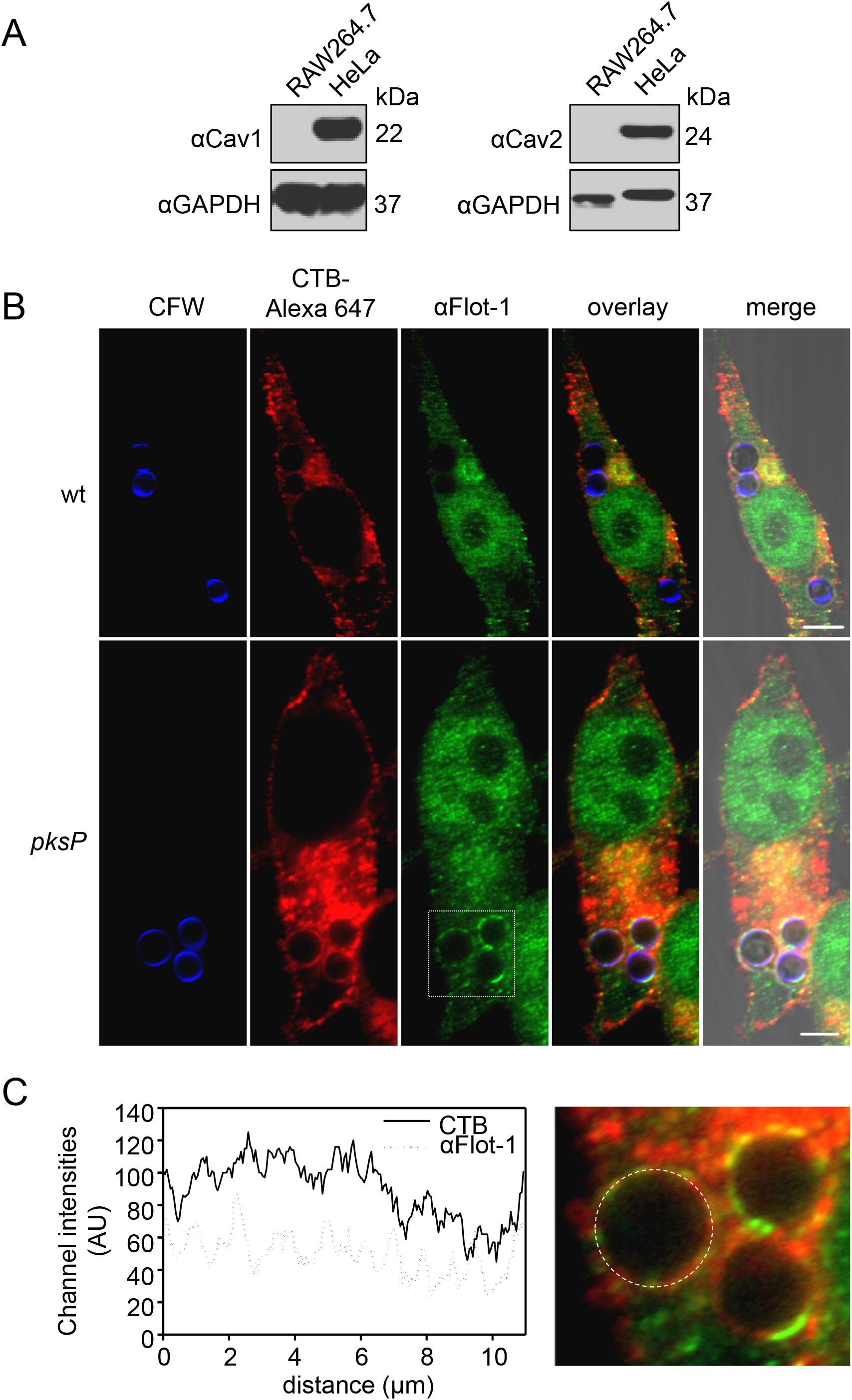
Different amounts of GM1 ganglioside and Flot-1 in conidia-containing PLs. **A)** Western blot analysis for detection of caveolin in RAW264.7 and HeLa cells. αGAPDH = control. **B)** Detection of Flot-1 and lipid rafts in conidia-containing phagolysosomal membrane. Conidia were stained with CFW (blue), RAW264.7 macrophages for lipid rafts and Flot-1 with CTB-Alexa 647 and an αFlot-1 antibody, respectively. Scale bar = 5 μm **C)** Channel intensity plot (left) reflecting the CTB and αFlot-1 fluorescence signal of the phagolysosomal membrane surrounding the *pksP* conidium (right).

### The number of *pksP* conidia residing in acidified PLs is drastically reduced in macrophages of *Flot-1/Flot-2* double knockout mice

In mice and humans, two flotillins are known, designated flotillin-1 (Flot-1) and flotillin-2 (Flot-2) (Babuke, Ruonala et al., 2009, Frick, Bright et al., 2007). Both isoforms are integrated in lipid rafts by *N*-terminal hairpin structures and together with other proteins are suggested to form hetero-oligomeric complexes (Solis, Hoegg et al., 2007). Knockdown of either *Flot-1* or *Flot-2* in the murine macrophage-like cell line J774A.1 reduced the acidification of PLs containing *pksP* conidia; the effect by knockdown of *Flot-1* was more pronounced (Fig. S5). Simultaneous knockdown of both *Flot-1* and *Flot-2* further decreased the percentage of acidified PLs containing wild-type conidia.

This initial finding was substantiated by the analysis of bone marrow-derived macrophages (BMDMs) from *Flot-1/Flot-2* double knockout (Flot^-/-^) mice (Bitsikas, Riento et al., 2014) (Fig. 4A). Compared to C57BL/6 wild-type mice, BMDMs of Flot^-/-^ mice had impaired acidification of PLs containing *pksP* conidia (Fig. 4B). As expected, melanized wild-type conidia containing PLs showed no significant acidification (Fig 4B).

**Figure 4.**
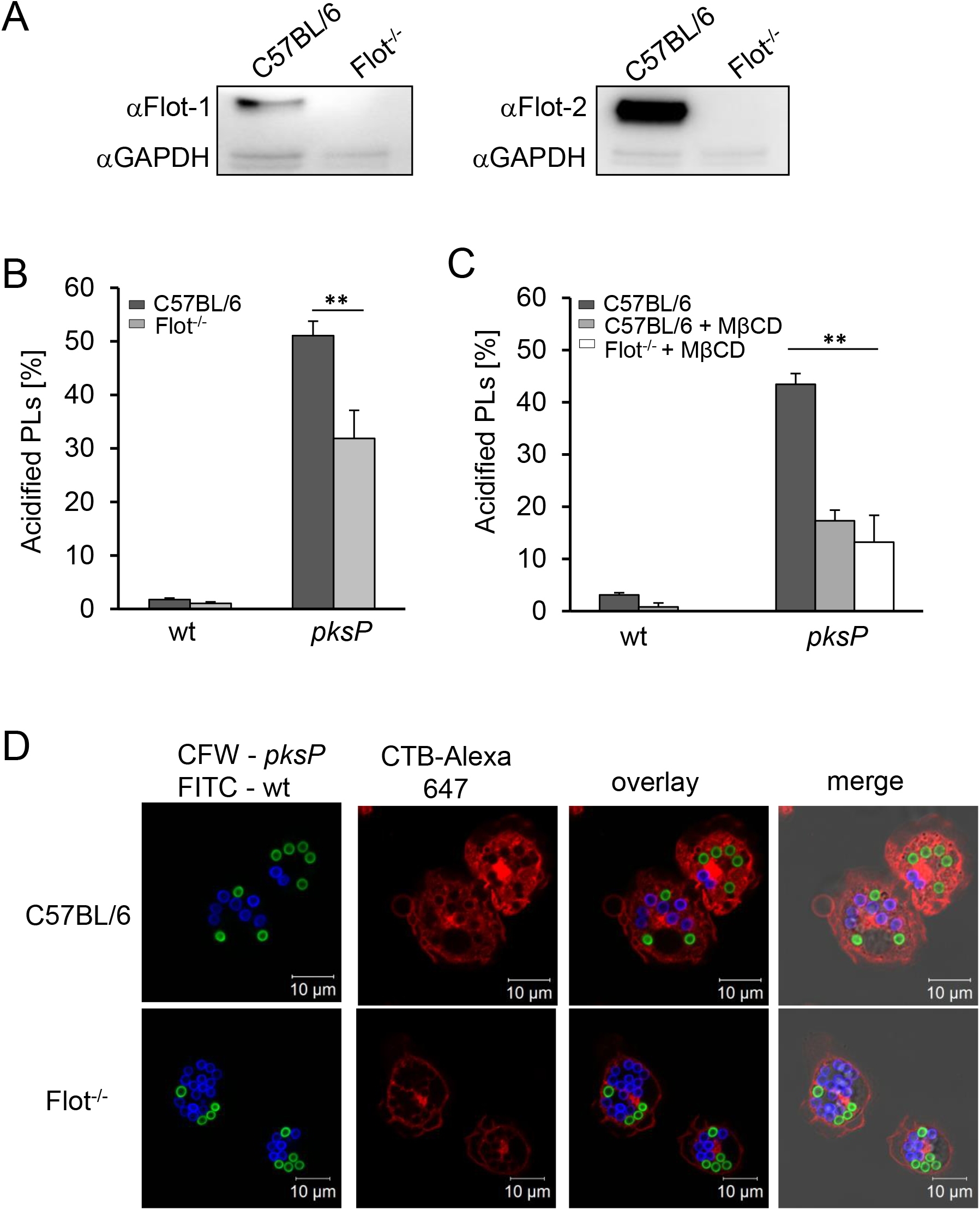
*A. fumigatus* wild-type conidia interfere with flotillin-dependent formation of phagolysosomal lipid rafts. **A)** Western blot analysis for detection of Flot-1 and Flot-2 in murine BMDMs of Flot^-/-^ and wildtype mice (C57BL/6). αGAPDH = control. **B)** Quantification of conidia-containing acidified PLs of wild-type or Flot^-/-^ BMDMs. Data are represented as mean ± SD; ** = *p* < 0.001. **C)** Quantification of acidified PLs in BMDMs from wild-type (C57BL/6) or Flot^-/-^ mice. When indicated, MβCD was added. Data are represented as mean ± SD; ** = *p* < 0.001. **D)** Coincubation of BMDMs with wild-type (FITC-labeled, green) and *pksP* (CFW-labeled, blue) conidia. CTB-Alexa 647 stained GM1 gangliosides (red). See also Figure S5.

We also confirmed that in primary mouse BMDMs acidification of *A. fumigatus* conidia-containing PLs was dependent on lipid-raft microdomains. In primary macrophages, only about 5% of PLs containing wild-type conidia were acidified, and this percentage was further reduced by treatment with MβCD. Furthermore, no phagolysosomal acidification was detectable in Flot^-/-^ macrophages treated with MβCD (Fig. 4C). The number of acidified PLs containing *pksP* conidia was drastically reduced by treatment with MβCD and was not further decreased in Flot^-/-^ macrophages treated with MβCD (Fig. 4C), indicating that flotillin and cholesterol function in the same pathway to enable phagolysosomal acidification. These results were further substantiated by using Flot^-/-^ knockout BMDMs which clearly showed reduced lipid-raft microdomains indicated by GM1 staining in the membrane of PLs containing either wild-type or *pksP* conidia (Fig. 4D).

### Assembly of vATPase and NADPH oxidase, and phagocytosis of conidia depend on flotillin-containing lipid-raft microdomains

Previous studies reported that in maturing phagosomes, vATPase is assembled in lipid-raft microdomains (Dermine, Duclos et al., 2001, Lafourcade, Sobo et al., 2008). To proof that flotillin as a lipid-raft chaperone is in fact required for this process, the assembly of vATPase in macrophages isolated from wild-type and Flot^-/-^ mice, was quantified (Fig. 5A). The percentage of phagolysosomal membranes showing a high degree of assembled vATPases when *pksP* conidia were ingested was reduced from 30 % in wild-type macrophages to about 10 % in Flot^-/-^ macrophages, indicating the dependency of vATPase assembly on flotillins, as earlier shown for cells of the murine macrophage-like cell line J774 (Dermine et al., 2001). STED (Stimulated Emission Depletion) microscopy confirmed colocalization of Flot-1 and vATPase V1 complex (Fig. 5B) at the phagolysosomal membrane.

**Figure 5.**
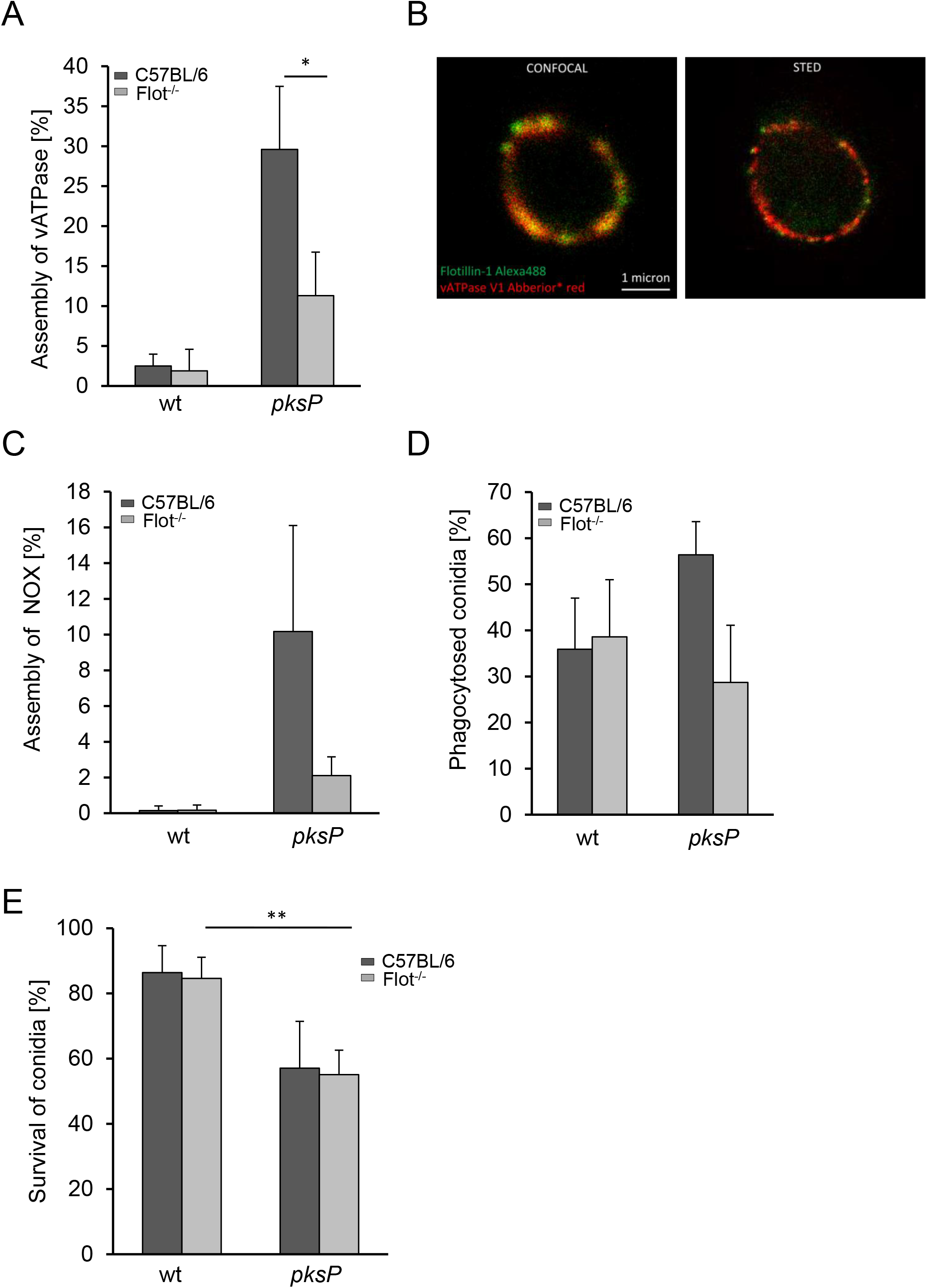
Flotillin-dependent lipid rafts are required for vATPase and NADPH oxidase assembly and phagocytosis of conidia. **A)** vATPase assembly in BMDMs isolated from C57BL/6 wild-type and Flot^-/-^ mice measured by immunofluorescence. Data are represented as mean ± SD; * = *p* < 0.05. **B)** Colocalization of vATPase and Flot-1. STED microscopy of isolated PLs stained with anti-flotillin-1 and anti-vATPase V_1_ antibody. C) NADPH oxidase assembly in BMDMs isolated from C57BL/6 wild-type and Flot^-/-^ mice measured by immunofluorescence. See also Figure S6. For vATPase and NADPH oxidase at least 100 intracellular conidia were evaluated for the presence of a fluorescence signal. The values represent mean ± SD of three independent experiments; **D)** Phagocytosis of conidia by Flot^-/-^ BMDMs. E) Killing of conidia by Flot^-/-^ BMDMs. The values represent mean ± SD of three independent experiments; ** = *p* < 0.001.

Interestingly, immunofluorescence revealed that the assembly of the NADPH oxidase complex on the phagolysosomal membrane was also affected by the absence of flotillins (Fig. S6). As shown in Fig. 5C, the percentage of PLs showing assembled NADPH oxidase complex when *pksP* conidia were ingested was reduced from 10 % in wild-type macrophages to 2 % in Flot^-/-^ BMDMs, also indicating the dependency of NADPH assembly on flotillins. Taken together, deletion of flotillin genes fully phenocopied the effect of melanin on PLs. As shown in Fig. 5D, phagocytosis rates of *pksP* conidia were reduced in Flot^-/-^ BMDMs. There was no effect on the phagocytosis rate of wild-type conidia, since these conidia *via* their DHN-melanin layer most likely already led to reduced flotillins in the membrane forming the phagocytic cup. Intracellular killing of conidia was not affected by Flot^-/-^ BMDMs compared to wild-type BMDMs (Fig. 5E).

### *FLOT1* SNP results in heightened susceptibility for invasive aspergillosis in hematopoietic stem-cell transplant recipients

Our data suggested that flotillin-dependent lipid-raft microdomain formation plays a key role in immunity to infection with *A. fumigatus*. Therefore, we next explored the role of this pathway in humans. We screened for single nucleotide polymorphisms (SNPs) and their association with the risk of invasive aspergillosis (IA) in a cohort of hematopoietic stem-cell transplant (HSCT) recipients (Stappers, Clark et al., 2018). Whereas for the *FLOT2* gene there was no SNP associated with risk of infection, remarkably, rs3094127, a SNP (T>C) in the last intron of the *FLOT1* gene that is lacking in the corresponding *FLOT1* gene in mice (Fig. 6A), was associated with an increased risk of IA after transplantation (Fig. 6B). The cumulative incidence of IA for donor rs309412 was 38.5% for CC (*p* = 0.04), 29.6% for TC (*p* = 0.05) and 19.6% for TT genotypes, respectively (Fig. 6B).

**Figure 6.**
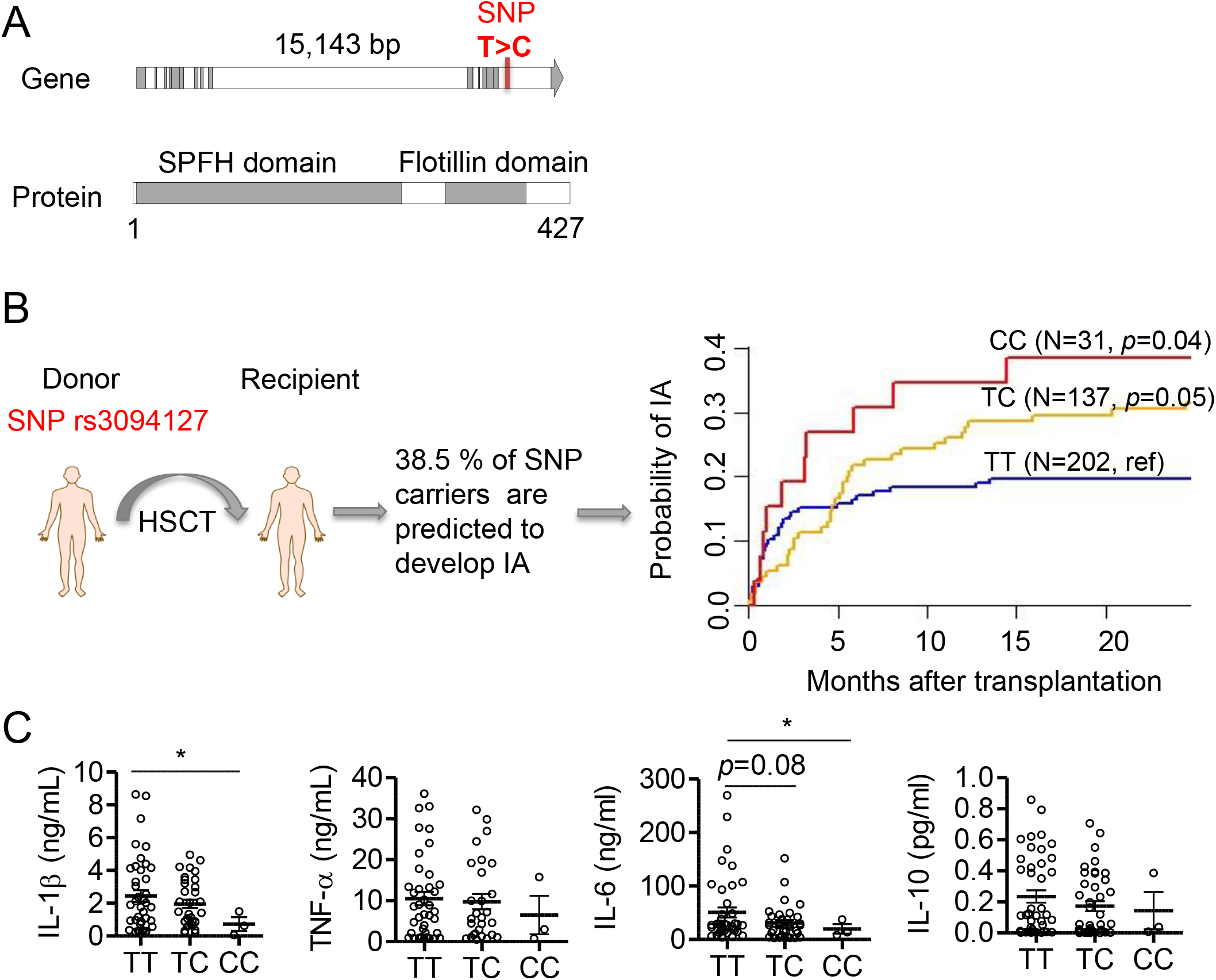
The rs3094127 SNP in *FLOT1* is associated with increased risk for invasive aspergillosis in a high-risk group of hematopoietic stem-cell recipients. A) Position of SNP found in human *FLOT1* gene. The SNP changes T to C. B) Cumulative incidence of IA in recipients of allogeneic hematopoietic stem-cell transplantation according to donor rs3094127 genotypes in *FLOT1*. Data were censored at 24 months, and relapse and death were competing events. *p* values are for Gray’s test. C) Cytokines released by monocyte-derived macrophages obtained from healthy donors with different *FLOT1* genotypes after stimulation with *A. fumigatus* conidia for 24 h.

In a multivariate model accounting for clinical variables associated with or tending towards IA in our cohort, the CC genotype at rs3094127 in *FLOT1* remained an independent predictor of IA (HR 1.89; 95 % CI 1.14-5.67; *p* = 0.03). The *FLOT1* genotypes had no impact in overall survival. To demonstrate a functional effect of the SNP in *Flot-1* on myeloid cell function, we analyzed the responses of monocyte-derived macrophages isolated from healthy genotyped donors. We found that macrophages from the individuals carrying this SNP produced significantly less IL-1β and IL-6 following *in vitro* stimulation with *A. fumigatus* conidia compared to controls, whereas cytokines TNF-α and IL10 did not show significant differences (Fig. 6C).

## DISCUSSION

Our study adds novel mechanistic insight on the role of lipid raft-based signaling in regulation of phagosome biogenesis and provides a new molecular virulence mechanism of melanin-induced inhibition of this process *via* the disruption of lipid rafts. The lipid microdomains (rafts) hypothesis was orginally proposed by Simons and Ikonen (Simons & Ikonen, 1997), imagining these lipid rafts as floating islands in the membrane (Luo, Wang et al., 2008, Triantafilou, Miyake et al., 2002). Lipid rafts are defined as small (10–200 nm) heterogeneous, highly dynamic, sterol (cholesterol), sphingolipid and protein-enriched domains that compartmentalize the cellular processes (Pike, 2006, Varshney, Yadav et al., 2016). One of the widely appreciated roles of lipid rafts is the recruitment and concentration of molecules involved in cellular signalling. The formation of a molecular cluster and their signal transduction machinery in membrane rafts leads to enhanced signalling efficiency (Triantafilou et al., 2002). It is thus not surprising that lipid rafts are required for immunity (Varshney et al., 2016). The exact role of lipid rafts and their composition are still a matter of debate. Nevertheless, the differentiation of membrane microdomains into sub-classes of lipid-raft domains and caveolae domains is widely accepted (Lingwood & Simons, 2010, Pike, 2009).

Here, we characterized lipid rafts in the phagolysosomal membrane by using a variety of marker molecules like GM1 ganglioside and cholesterol. To further specify the membrane microdomains, we assessed the localization of typical lipid raft marker proteins during PL maturation. Our findings are in accordance with studies underlining that caveolins are not coexpressed in murine macrophages (Gargalovic & Dory, 2001, Nagao, Ishii et al., 2010). Instead, our data indicate that flotillin-dependent lipid-raft membrane domains play a pivotal role in phagosome biogenesis and acidification of PLs (Lafourcade et al., 2008) and that these membrane structures are targeted by *A. fumigatus* conidia to interfere with the maturation of this compartment. These findings were confirmed by the observation that flotillin-1, a marker for a different subset of lipid rafts and assumed to act as scaffolding protein that stabilizes lipid rafts (Banning et al., 2011, Otto & Nichols, 2011, Stuermer, 2011), colocalized with GM1 ganglioside on phagolysosomal membranes of RAW264.7 macrophages containing *pksP* conidia but not melanized conidia. Further, knockdown and knockout of the flotillin genes had a major influence on the acidification of PLs and the lipid-raft assembly on the phagolysosomal membrane. Consistently, flotillins were found to be required for vATPase assembly, explaining the lack of acidification of PLs containing wild-type conidia.

Flotillins belong to a family of lipid-raft-associated integral membrane proteins. Flotillin members are ubiquitously expressed and located to non-caveolar microdomains on the cell membrane. Two flotillin members have been described, flotillin-1 and flotillin-2 (Otto & Nichols, 2011, Vieira, Correa et al., 2010). They constitutively associate with lipid rafts by acylation (a single palmitate in flotillin-1, a myristate and three palmitates in flotillin-2), oligomerization and cholesterol binding (Meister & Tikkanen, 2014). Flotillins have been long considered markers of lipid rafts because they are detergent insoluble and float in sucrose density gradients. Consequently, flotillins, either on their own or in combination with the respective other flotillin, have been implicated in numerous signalling events and pathways that are thought to be organised in lipid rafts (Babuke & Tikkanen, 2007). As shown here, by emplyoing immunostaining, BMDMs of Flot^-/-^ mice and knockdown cell lines, the lipid rafts on the phagolysosomal membrane contain and require flotillin proteins.

An important aspect found here is that flotillin-dependent lipid-rafts form a platform for vATPase assembly and NADPH oxidase complex assembly on the phagolysosomal membrane. Further, they contribute to phagocytosis. These findings not only resulted from experiments with knockout cells but also from the colocalization of vATPase and Flot-1 on the phagolysosomal membrane. The vATPase is required for pumping protons in the lumen of the PL and thereby acidifying its content. The V_1_ complex of vATPase is located in the cytoplasm and contains the active site responsible for hydrolyzing ATP. The reversible binding of the V_1_ to the V_0_ complex in the phagosomal membrane regulates the activity of vATPase (Cotter et al., 2015). Little is known about the role of the vATPase and its regulation associated with the endocytic pathways of immune cells although this multiprotein complex is crucial for the antimicrobial properties of professional phagocytes, *e.g*., intracellular killing, digestion, and presentation of antigenic epitopes (Cotter et al., 2015, Lukacs, Rotstein et al., 1990). Genome wide knockout studies in *Saccharomyces cerevisiae* showed that the presence of sphingolipids strongly influences the assembly of the vATPase complex (Chung, Lester et al., 2003, Finnigan, Ryan et al., 2011). In addition to cholesterol, sphingolipids are key components of lipid rafts (Fessler & Parks, 2011, Pike, 2009, Simons & Gerl, 2010, Simons & Toomre, 2000). There is a direct link between lipidraft microdomains and regulation of the vATPase (Dhungana, Merrick et al., 2009, Foster, De Hoog et al., 2003, Lafourcade et al., 2008). Here, by using knockout BMDMs and knockdown cell lines we obtained evidence that the assembly of a functional vATPase on the phagolysosomal membrane requires flotillin-dependent lipid rafts. In line, by coinfecting macrophages with both wild-type and *pksP* conidia, not only different acidification patterns of PLs in the same cell occurred but also different localization patterns of the cytoplasmic V1 vATPase complex. All these effects were due to different amounts of lipid rafts in the phagolysosomal membrane. Thus, dysregulation of phagolysosomal lipid rafts by melanized conidia is not an overall mechanism of the entire cell but locally restricted to the specific phagolysosomal compartment containing a conidium. These findings correspond with previous results showing that during PL maturation the association of phagolysosomal compartments with lipid rafts governs increased acidification due to higher vATPase assembly rates (Lafourcade et al., 2008). As shown for LAP (Kyrmizi et al., 2018), the restriction of these effects to a distinct PL is most likely due to the binding of activated calmodulin to the phagolysosomal membrane. If there is free Ca^2+^ available for release from the phagolysosomal lumen to the cytosolic part of the phagosomal membrane, calmodulin on the cytosolic part of the membrane is activated and as a result the PL is functional, as seen here for PLs containing *pksP* conidia.

An interesting finding is the dependence of NADPH oxidase complex assembly on flotillin-dependent lipid rafts. It is conceivable that defects in flotillins result in immune suppression, *e.g*., as shown here phagocytosis was reduced in Flot^-/-^ BMDMs or reactive oxygen species (ROS) produced by NADPH oxidase are required for induction of LAP (Martinez, Malireddi et al., 2015). This assumption is supported by the reduced phagocytosis rate of *pksP* conidia in Flot^-/-^ BMDMs suggesting that lipid rafts are required as signaling platforms for recognition of conidia. By contrast, wild-type conidia did not show different phagocytosis rates because these conidia *via* their DHN-melanin layer most likely already disturbed flotillins in the membrane forming the phagocytic cup. In line, recently it was shown that flotillin-1 facilitates inflammatory toll-like receptor 3 signaling in human endothelial cells (Fork, Hitzel et al., 2014).

Until now, the regulation of lipid-raft formation is a matter of debate. Here, we found that inhibition of Ca^2+^-dependent calmodulin activity or melanin-dependent Ca^2+^ sequestration in the PL reduced the presence of lipid rafts on the phagolysosomal membrane. Calmodulin is a versatile Ca^2+^-sensor/transducer protein that modulates hundreds of enzymes, channels, transport systems, transcription factors, adaptors and other structural proteins, controlling in this manner multiple cellular functions. It can regulate target proteins in a Ca^2+^-dependent and Ca^2+^-independent manner (Villalobo, 2018). Since we found clear differences in the concentration of free Ca^2+^ in PLs containing pigmentless *pksP* conidia *versus* melanized conidia, this data suggests that the available free Ca^2+^ ions for calmodulin are reduced and thereby calmodulin activation and lipid-raft formation. This conclusion was substantiated by the finding of reduced lipid-raft formation on the phagolysosomal membrane when cells were treated with a calmodulin inhibitor. Our data well agree with our previous observation that early, transient and selective calmodulin localization was exclusively observed in PLs of melanin-deficient conidia (Kyrmizi et al., 2018).

Until now, for different pathogens such as viruses, bacteria and protozoa, it was shown that they can use host-cell lipid rafts to secure their entrance and maintenance inside target cells (Manes, del Real et al., 2003, Vieira et al., 2010). Different viruses have evolved strategies to subvert raft-associated signalling, enabling their efficient replication in immune cells, and at the same time blocking the immune response that is elicited by the target cells (Hawkes & Mak, 2006). Likewise, several bacteria interact with host lipid rafts to enter and survive inside the cell (Hawkes & Mak, 2006, Manes et al., 2003). The mechanisms that underlie this interaction are starting to be unravelled. Activation of secretion, binding, perforation of the host-cell membrane and signalling to trigger bacterial phagocytosis are involved with components of membrane microdomains (Lafont & van der Goot, 2005, Vieira et al., 2010).

Maza and colleagues (2008) investigated yeast forms of the fungal pathogen *Paracoccidioides brasiliensis* in the context of kinase signalling. This pathogen apparently promotes the aggregation of lipid rafts in epithelial cellls allowing fungal adhesion (Maza, Straus et al., 2008). Here, we uncovered a novel virulence mechanism. Unlike many facultative intracellular pathogens, as shown here *A. fumigatus* evades phagocytes by impairing lipid-raft formation in phagolysosomal membranes most likely *via* Ca^2+^ sequestration by melanized conidia. Thereby, melanized conidia prevent lipid raft-associated assembly of the signal platforms required, *e.g*., for assembly of vATPase and NADPH oxidase. However, alternatively it is also conceivable that incomplete or only partial fusions have occurred between endosomal and lysosomal vesicles instead of complete fusions with entire mixing of the fusion partners’ membranes (Desjardins, 1995, Haas, 2007). Since lipid rafts are also associated with functional membrane trafficking (Simons & Gerl, 2010), missing microdomains in the wild-type conidia-containing phagolysosomal membranes could be responsible for the lack of integration of lysosomal membrane constituents, which then leads to PLs devoid of lipid-raft microdomains.

Recently, we reported that DHN-melanin inhibits the activation of a noncanonical LAP autophagy pathway (Akoumianaki et al., 2016), which is regulated by NADPH oxidase and which promotes fungal killing. It was found that DHN-melanin inhibits NADPH oxidase-dependent activation of LAP by excluding the p22phox subunit from the PL. Also, NOX2-generated ROS are necessary for LC3 recruitment to phagosomes (Huang, Canadien et al., 2009). Here, we provide a model explaining that conidial melanin interferes with lipid-raft formation that is required for NADPH oxidase assembly and most likely all further processes assigned to the activity of melanin.

The importance of the lipid-raft component flotillin for pathogenicity was impressively demonstrated by the analysis of a total of 370 hematologic patients undergoing allogeneic hematopoietic stem-cell transplantation. We identified a SNP in a region of the *FLOT1* gene that is not present in mice and which results in heightened susceptibility for invasive aspergillosis. Macrophages of individuals homozygous for this SNP showed reduced extracellular amounts of IL-1ß and IL-6. IL-1ß has been clearly shown to be required for defense against *A. fumigatus* infection (Sainz, Perez et al., 2008, Wojtowicz, Gresnigt et al., 2015) and flotillin dependent lipid rafts have been shown to affect cytokine secretion (Kay, Murray et al., 2006). These data show the functional impairment caused by the SNP and the importance of Flot-1-dependent lipid rafts for immunity against infections. In the *FLOT2*, gene we did not detect a SNP that could be correlated with susceptibility for invasive aspergillosis. This finding is also supported by the observation that the knockdown of either *FLOT1* or *FLOT2* in the murine macrophage cell line J774A.1 both reduced the acidification of PLs containing *pksP* conidia, but knockdown of *FLOT1* had a greater effect. Until now, it is a matter of debate whether flotillins 1 and 2 are codependent in their cellular functions, or whether they can also function individually that was shown for a number of cellular effects (Bitsikas et al., 2014, Langhorst, Reuter et al., 2008, Langhorst, Solis et al., 2007, Neumann-Giesen, Fernow et al., 2007, Schneider, Rajendran et al., 2008, Stuermer, 2011, Vieira et al., 2010). Thus, it is likely that flotillins can act as heteromers but also independent of each other.

Collectively, our data provide new insight into the importance of lipid rafts for immunity against human pathogenic fungi and as a target for pathogens. Our study adds novel mechanistic insight into the regulation and formation of lipid rafts, elucidates a role for lipid raft-based signaling in regulation of phagosome biogenesis and reports a new molecular virulence mechanism *via* the disruption of lipid-raft microdomains.

Molecules on the fungal surface and excreted molecules belong to the first structures interacting with host cells. Although the recognition of immunological cell wall structures like β-1,3-glucans (Steele, Rapaka et al., 2005) which are more susceptible for receptors on *pksP* conidia (Luther et al., 2007) leads to an activation of immune cells by generation of an inflammatory response through production of chemokines and cytokines (Chai et al., 2010), it does not necessarily result in a higher activation of the endocytic pathway with accompanied acidification of phagolysosomal compartments. This was concluded here from experiments, in which the same macrophage had phagocytosed both a wild-type and a *pksP* conidium, which, in the same cell, ended in a neutral and acidified phagolysosome, respectively. Instead, each ingested particle determines its intracellular fate *via* influencing the endocytic pathway by its morphological and chemical properties, in the case of *A. fumigatus* conidia by the presence or absence of the DHN-melanin layer. As shown here, abolishing the phagolysosomal acidification by disturbance of vATPase assembly resulted in increased host cell damage, interestingly to the same extent for both wild-type and *pksP* conidia. Thus, phagolysosomal acidification strongly contributes to the effective killing of conidia (Jahn, Langfelder et al., 2002) and thereby protects to a certain extent the host cell from outgrowth of the ingested pathogen. Inhibition of phagolysosomal acidification and reduction of NADPH oxidase complex assembly by DHN-melanin allows wild-type conidia to create a favorable niche with a nearly neutral pH, which prevents activation of hydrolytic enzymes and thereby phagolysosomal digestion.

## EXPERIMENTAL PROCEDURES

### Cultivation of fungal strains, cell lines and infection experiments

The *A. fumigatus* strains used in this study, the ATCC46645 wild type and the non-pigmented *pksP* mutant were cultivated on *Aspergillus* minimal medium (AMM) agar plates as described elsewhere (Gsaller, Hortschansky et al., 2014, Jahn et al., 1997). MH-S (ATCC:CRL-2019), J774A.1 (ATCC:TIB-67), HeLa (ATCC:CCL-2), and RAW264.7 cells (ATCC:TIB-71) were cultivated in RPMI 1640 and DMEM, respectively, at 37 °C, 5 % (v/v) CO_2_ in a humidified chamber. A detailed protocol is provided in the Supplemental Information. The calmodulin inhibitor W7 (A3281, Sigma-Aldrich) was used in a concentration of 10 μM.

### Host cell damage assay

Host cell damage caused by *A. fumigatus* was measured by the previously described ^51^Cr release assay (Filler, Swerdloff et al., 1995). A detailed protocol is provided in the Supplemental Information.

### Immunofluorescence and sample preparation for CLSM and STED

See the Supplemental Information for detailed description.

### Quantitation of phagolysosomal acidification and lipid raft recruitment

Prior to infection, macrophages were preloaded with 50 nM LysoTracker Red DND-99 (Life Technologies) in medium for 1 h in the absence or presence of 7.5 mM methyl-β-cyclodextrin (MβCD) (Sigma-Aldrich) or after addition of 1 μM Alexa Fluor 647-conjugated Cholera Toxin B (Life Technologies), cells were infected with *A. fumigatus* conidia. Then, fixed or live cells were subjected to microscopic analysis. The values represent mean ± SD of three different experiments. A detailed protocol is provided in the Supplemental Information.

### Subcellular fractionation and Western blot analysis

After coincubation of conidia and immune cells, cytoplasmic and host membrane-containing cell fractions were obtained using the Qproteome Cell Compartment Kit (Qiagen) according to the manufacturer’s instructions. PLs were prepared from RAW264.7 macrophages as previously described (Akoumianaki et al., 2016). A detailed protocol is provided in the Supplemental Information.

### Extraction and quantification of cholesterol

Cholesterol measurements were performed using liquid chromatography coupled to triple-quadrupole mass spectrometry (LC-MS/MS). A detailed protocol is provided in the Supplemental Information.

### Calcium staining

Cells were loaded at a 1:1 ratio of Fluo-4 AM (Life Technologies) and cell medium (DMEM) 30 min before coincubation. After infection with conidia (MOI = 5) cells were coincubated for 2 h and PLs were isolated as described before.

### Knockdown of Flotillin genes

J774A.1 cells were seeded with a density of 3×10^5^ or 1×10^5^ in 6 or 24 well plates in DMEM supplemented with 10 % (v/v) FBS and 1 % (w/v) ultraglutamine. After 24 hours the cells were transfected with 40 nM target specific siRNA (Santa Cruz Biotechnology) and 3-6 μl Lipofectamin 2000 (Life Technologies). See the Supplemental Information for detailed description.

### Mouse strains and isolation of murine bone marrow-derived macrophages

Flotillin-1, flotillin-2 (*Flot-1, Flot-2*) double knockout (Flot^-/-^) mice (Bitsikas et al., 2014) were a kind gift of Ben J. Nichols (Cambridge, UK). Bone marrow cells were harvested from femurs and tibias of specific pathogen-free mice according to a procedure described elsewhere (Zhang, Goncalves et al., 2001). All animals were cared for in accordance with the European animal welfare regulation. A detailed protocol is provided in the Supplemental Information.

### Human studies

A total of 370 hematologic patients undergoing allogeneic hematopoietic stem-cell transplantation at the Hospital of Santa Maria, Lisbon and Instituto Português de Oncologia (IPO), Porto, between 2009 and 2014 were enrolled in the study (Stappers et al., 2018), including 79 cases of probable/proven aspergillosis and 244 uninfected controls. The cases of invasive aspergillosis were identified and classified as ‘probable’ or ‘proven’ according to the revised standard criteria from the European Organization for Research and Treatment of Cancer/Mycology Study Group (EORTC/MSG) (De Pauw, Walsh et al., 2008). Study approval for the genetic association study was obtained from the Ethics Subcommittee for Life and Health Sciences of the University of Minho, Portugal (125/014), the Ethics Committee for Health of the Instituto Português de Oncología - Porto, Portugal (26/015), the Ethics Committee of the Lisbon Academic Medical Center, Portugal (632/014), and the National Commission for the Protection of Data, Portugal (1950/015). Approval for the functional studies on human cells was provided by Ethics Subcommittee for Life and Health Sciences of the University of Minho, Portugal (SECVS-014/2015). All individuals provided written informed consent in accordance with the Declaration of Helsinki. Genomic DNA was isolated from whole blood using the QIAcube automated system (Qiagen). Genotyping of the rs3094127 SNP in the *FLOT1* gene was performed using KASPar assays (LGC Genomics) in an Applied Biosystems 7500 Fast PCR system (Thermo Fisher).

### Statistical analysis

Data are expressed as mean ± SD. *P* values were calculated by a one-way ANOVA (Bonferroni’s post hoc test). For single comparison, *p* values were calculated by a two-tailed Student’s t test.

## EXPANDED VIEW CONTENT

Expanded View Content includes detailed experimental procedures, six supplemental figures and two supplemental movies.

## AUTHOR CONTRIBUTIONS

F.S., A.T., M.R., Z.C., H.S., S.G., A.C., M.H.G., and T.H. conducted experiments and analyzed data, A.A.B., M.T.F., G.C., C.C., A.C., J.F.L., A.C.Jr., C.E., and S.G.F. designed research and analyzed data, F.S., H.S., A.T., M.R., T.H., G.C., S.G.F. and A.A.B. wrote the paper.

## ACKNOWLEDGEMENTS

We are extremely grateful to Ben J. Nichols (Cambridge, UK) for providing flotillin knockout mice. We thank Maria Straßburger for projecting mouse breeding and Anna Runtze and Muhammad Rafiq for initial experiments. We thank Matthew Blango for critically reading the manuscript. This work was supported by the excellence graduate school Jena School for Microbial Communication (JSMC) funded by the Deutsche Forschungsgemeinschaft (DFG), the International Leibniz Research School (ILRS) as part of the JSMC, the Leibniz Science Campus ‘InfectoOptics’ and the DFG-funded CRC/TR 124 ‘Pathogenic fungi and their human host - Networks of interaction - FungiNet’ (project A1 to A.A.B. and project B4 to M.T.F.) and CRC 1278 ‘Polymer-based nanoparticle libraries for targeted anti-inflammatory strategies - PolyTarget’ (project B02 to A.A.B. and Z01 to M.T.F.). C.C. and A.C were supported by the Northern Portugal Regional Operational Programme (NORTE 2020), under the Portugal 2020 Partnership Agreement, through the European Regional Development Fund (FEDER) (NORTE-01-0145-FEDER-000013), and by the Fundação para a Ciência e Tecnologia (FCT) (SFRH/BPD/96176/2013 to C.C. and IF/00735/2014 to A.C.). There is no conflict of interest for any of the authors.

## REFERENCES

Akoumianaki T, Kyrmizi I, Valsecchi I, Gresnigt MS, Samonis G, Drakos E, Boumpas D, Muszkieta L, Prevost MC, Kontoyiannis DP, Chavakis T, Netea MG, van de Veerdonk FL, Brakhage AA, El-Benna J, Beauvais A, Latge JP, Chamilos G (2016) *Aspergillus* cell wall melanin blocks LC3-associated phagocytosis to promote pathogenicity. Cell Host Microbe 19: 79–90

Alvarez M, Casadevall A (2006) Phagosome extrusion and host-cell survival after *Cryptococcus neoformans* phagocytosis by macrophages. Curr Biol 16: 2161–5

Andrianaki AM, Kyrmizi I, Thanopoulou K, Baldin C, Drakos E, Soliman SSM, Shetty AC, McCracken C, Akoumianaki T, Stylianou K, Ioannou P, Pontikoglou C, Papadaki HA, Tzardi M, Belle V, Etienne E, Beauvais A, Samonis G, Kontoyiannis DP, Andreakos E et al. (2018) Iron restriction inside macrophages regulates pulmonary host defense against *Rhizopus* species. Nat Commun 9: 3333

Babuke T, Ruonala M, Meister M, Amaddii M, Genzler C, Esposito A, Tikkanen R (2009) Heterooligomerization of reggie-1/flotillin-2 and reggie-2/flotillin-1 is required for their endocytosis. Cell Signal 21: 1287–97

Babuke T, Tikkanen R (2007) Dissecting the molecular function of reggie/flotillin proteins. Eur J Cell Biol 86: 525–32

Banning A, Tomasovic A, Tikkanen R (2011) Functional aspects of membrane association of reggie/flotillin proteins. Curr Protein Pept Sci 12: 725–35

Batanghari JW, Deepe GS, Jr., Di Cera E, Goldman WE (1998) *Histoplasma* acquisition of calcium and expression of CBP1 during intracellular parasitism. Mol Microbiol 27: 531–9

Bitsikas V, Riento K, Howe JD, Barry NP, Nichols BJ (2014) The role of flotillins in regulating abeta production, investigated using flotillin 1-/-, flotillin 2-/- double knockout mice. PLoS One 9: e85217

Brown GD, Denning DW, Gow NA, Levitz SM, Netea MG, White TC (2012) Hidden killers: human fungal infections. Sci Transl Med 4: 165rv13

Cambier CJ, Falkow S, Ramakrishnan L (2014) Host evasion and exploitation schemes of *Mycobacterium tuberculosis*. Cell 159: 1497–509

Carvalho F, Sousa S, Cabanes D (2014) How *Listeria monocytogenes* organizes its surface for virulence. Front Cell Infect Microbiol 4: 48

Chai LY, Netea MG, Sugui J, Vonk AG, van de Sande WW, Warris A, Kwon-Chung KJ, Kullberg BJ (2010) *Aspergillus fumigatus* conidial melanin modulates host cytokine response. Immunobiology 215: 915–20

Chung JH, Lester RL, Dickson RC (2003) Sphingolipid requirement for generation of a functional v1 component of the vacuolar ATPase. The Journal of biological chemistry 278: 28872–81

Cotter K, Stransky L, McGuire C, Forgac M (2015) Recent insights into the structure, regulation, and function of the V-ATPases. Trends Biochem Sci 40: 611–22

De Pauw B, Walsh TJ, Donnelly JP, Stevens DA, Edwards JE, Calandra T, Pappas PG, Maertens J, Lortholary O, Kauffman CA, Denning DW, Patterson TF, Maschmeyer G, Bille J, Dismukes WE, Herbrecht R, Hope WW, Kibbler CC, Kullberg BJ, Marr KA et al. (2008) Revised definitions of invasive fungal disease from the European Organization for Research and Treatment of Cancer/Invasive Fungal Infections Cooperative Group and the National Institute of Allergy and Infectious Diseases Mycoses Study Group (EORTC/MSG) Consensus Group. Clin Infect Dis 46: 1813–21

Dermine JF, Duclos S, Garin J, St-Louis F, Rea S, Parton RG, Desjardins M (2001) Flotillin-1-enriched lipid raft domains accumulate on maturing phagosomes. The Journal of biological chemistry 276: 18507–12

Desjardins M (1995) Biogenesis of phagolysosomes: the ‘kiss and run’ hypothesis. Trends Cell Biol 5: 183–6

Dhungana S, Merrick BA, Tomer KB, Fessler MB (2009) Quantitative proteomics analysis of macrophage rafts reveals compartmentalized activation of the proteasome and of proteasome-mediated ERK activation in response to lipopolysaccharide. Mol Cell Proteomics 8: 201–13

Ensminger AW (2016) *Legionella pneumophila,* armed to the hilt: justifying the largest arsenal of effectors in the bacterial world. Curr Opin Microbiol 29: 74–80

Fernandez-Arenas E, Bleck CK, Nombela C, Gil C, Griffiths G, Diez-Orejas R (2009) *Candida albicans* actively modulates intracellular membrane trafficking in mouse macrophage phagosomes. Cell Microbiol 11: 560–89

Fessler MB, Parks JS (2011) Intracellular lipid flux and membrane microdomains as organizing principles in inflammatory cell signaling. J Immunol 187: 1529–35

Filler SG, Swerdloff JN, Hobbs C, Luckett PM (1995) Penetration and damage of endothelial cells by *Candida albicans*. Infect Immun 63: 976–83

Finnigan GC, Ryan M, Stevens TH (2011) A genome-wide enhancer screen implicates sphingolipid composition in vacuolar ATPase function in *Saccharomyces cerevisiae*. Genetics 187: 771–83

Fork C, Hitzel J, Nichols BJ, Tikkanen R, Brandes RP (2014) Flotillin-1 facilitates toll-like receptor 3 signaling in human endothelial cells. Basic Res Cardiol 109: 439

Foster LJ, De Hoog CL, Mann M (2003) Unbiased quantitative proteomics of lipid rafts reveals high specificity for signaling factors. Proc Natl Acad Sci U S A 100: 5813–8

Frick M, Bright NA, Riento K, Bray A, Merrified C, Nichols BJ (2007) Coassembly of flotillins induces formation of membrane microdomains, membrane curvature, and vesicle budding. Curr Biol 17: 1151–6

Gargalovic P, Dory L (2001) Caveolin-1 and caveolin-2 expression in mouse macrophages. High density lipoprotein 3-stimulated secretion and a lack of significant subcellular co-localization. The Journal of biological chemistry 276: 26164–70

Gsaller F, Hortschansky P, Beattie SR, Klammer V, Tuppatsch K, Lechner BE, Rietzschel N, Werner ER, Vogan AA, Chung D, Muhlenhoff U, Kato M, Cramer RA, Brakhage AA, Haas H (2014) The Janus transcription factor HapX controls fungal adaptation to both iron starvation and iron excess. The EMBO journal 33: 2261–76

Haas A (2007) The phagosome: compartment with a license to kill. Traffic 8: 311–30

Hawkes DJ, Mak J (2006) Lipid membrane; a novel target for viral and bacterial pathogens. Curr Drug Targets 7: 1615–21

Heinekamp T, Thywissen A, Macheleidt J, Keller S, Valiante V, Brakhage AA (2012) *Aspergillus fumigatus* melanins: interference with the host endocytosis pathway and impact on virulence. Front Microbiol 3: 440

Huang J, Canadien V, Lam GY, Steinberg BE, Dinauer MC, Magalhaes MA, Glogauer M, Grinstein S, Brumell JH (2009) Activation of antibacterial autophagy by NADPH oxidases. Proc Natl Acad Sci U S A 106: 6226–31

Jahn B, Koch A, Schmidt A, Wanner G, Gehringer H, Bhakdi S, Brakhage AA (1997) Isolation and characterization of a pigmentless-conidium mutant of *Aspergillus fumigatus* with altered conidial surface and reduced virulence. Infect Immun 65: 5110–7

Jahn B, Langfelder K, Schneider U, Schindel C, Brakhage AA (2002) PKSP-dependent reduction of phagolysosome fusion and intracellular kill of *Aspergillus fumigatus* conidia by human monocyte-derived macrophages. Cell Microbiol 4: 793–803

Kay JG, Murray RZ, Pagan JK, Stow JL (2006) Cytokine secretion via cholesterol-rich lipid raft-associated SNAREs at the phagocytic cup. The Journal of biological chemistry 281: 11949–54

Keller P, Simons K (1998) Cholesterol is required for surface transport of influenza virus hemagglutinin. J Cell Biol 140: 1357–67

Kosmidis C, Denning DW (2015) The clinical spectrum of pulmonary aspergillosis. Thorax 70: 270–7

Kyrmizi I, Ferreira H, Carvalho A, Figueroa JAL, Zarmpas P, Cunha C, Akoumianaki T, Stylianou K, Deepe GS, Jr., Samonis G, Lacerda JF, Campos A, Jr., Kontoyiannis DP, Mihalopoulos N, Kwon-Chung KJ, El-Benna J, Valsecchi I, Beauvais A, Brakhage AA, Neves NM et al. (2018) Calcium sequestration by fungal melanin inhibits calcium-calmodulin signalling to prevent LC3-associated phagocytosis. Nat Microbiol 3: 791–803

Lafont F, van der Goot FG (2005) Bacterial invasion via lipid rafts. Cell Microbiol 7: 613–20

Lafourcade C, Sobo K, Kieffer-Jaquinod S, Garin J, van der Goot FG (2008) Regulation of the V-ATPase along the endocytic pathway occurs through reversible subunit association and membrane localization. PLoS One 3: e2758

Langfelder K, Streibel M, Jahn B, Haase G, Brakhage AA (2003) Biosynthesis of fungal melanins and their importance for human pathogenic fungi. Fungal Genet Biol 38: 143–58

Langhorst MF, Reuter A, Jaeger FA, Wippich FM, Luxenhofer G, Plattner H, Stuermer CA (2008) Trafficking of the microdomain scaffolding protein reggie-1/flotillin-2. Eur J Cell Biol 87: 211–26

Langhorst MF, Solis GP, Hannbeck S, Plattner H, Stuermer CA (2007) Linking membrane microdomains to the cytoskeleton: regulation of the lateral mobility of reggie-1/flotillin-2 by interaction with actin. FEBS Lett 581: 4697–703

Ledeen RW, Wu G (2015) The multi-tasked life of GM1 ganglioside, a true factotum of nature. Trends Biochem Sci 40: 407–18

Levitz SM, Nong SH, Seetoo KF, Harrison TS, Speizer RA, Simons ER (1999) *Cryptococcus neoformans* resides in an acidic phagolysosome of human macrophages. Infect Immun 67: 885–90

Lingwood D, Simons K (2010) Lipid rafts as a membrane-organizing principle. Science 327: 46–50

Lukacs GL, Rotstein OD, Grinstein S (1990) Phagosomal acidification is mediated by a vacuolar-type H(+)-ATPase in murine macrophages. The Journal of biological chemistry 265: 21099–107

Luo C, Wang K, Liu DQ, Li Y, Zhao QS (2008) The functional roles of lipid rafts in T cell activation, immune diseases and HIV infection and prevention. Cell Mol Immunol 5: 1–7

Luther K, Torosantucci A, Brakhage AA, Heesemann J, Ebel F (2007) Phagocytosis of *Aspergillus fumigatus* conidia by murine macrophages involves recognition by the dectin-1 beta-glucan receptor and Toll-like receptor 2. Cell Microbiol 9: 368–81

Ma H, Croudace JE, Lammas DA, May RC (2006) Expulsion of live pathogenic yeast by macrophages. Curr Biol 16: 2156–60

Manes S, del Real G, Martinez AC (2003) Pathogens: raft hijackers. Nat Rev Immunol 3: 557–68

Martinez J, Malireddi RK, Lu Q, Cunha LD, Pelletier S, Gingras S, Orchard R, Guan JL, Tan H, Peng J, Kanneganti TD, Virgin HW, Green DR (2015) Molecular characterization of LC3-associated phagocytosis reveals distinct roles for Rubicon, NOX2 and autophagy proteins. Nature cell biology 17: 893–906

Maza PK, Straus AH, Toledo MS, Takahashi HK, Suzuki E (2008) Interaction of epithelial cell membrane rafts with *Paracoccidioides brasiliensis* leads to fungal adhesion and Src-family kinase activation. Microbes Infect 10: 540–7

Mech F, Thywissen A, Guthke R, Brakhage AA, Figge MT (2011) Automated image analysis of the host-pathogen interaction between phagocytes and *Aspergillus fumigatus*. PLoS One 6: e19591

Meister M, Tikkanen R (2014) Endocytic trafficking of membrane-bound cargo: a flotillin point of view. Membranes (Basel) 4: 356–71

Nagao G, Ishii K, Hirota K, Makino K, Terada H (2010) Role of lipid rafts in phagocytic uptake of polystyrene latex microspheres by macrophages. Anticancer Res 30: 3167–76

Neumann-Giesen C, Fernow I, Amaddii M, Tikkanen R (2007) Role of EGF-induced tyrosine phosphorylation of reggie-1/flotillin-2 in cell spreading and signaling to the actin cytoskeleton. J Cell Sci 120: 395–406

Newman SL, Gootee L, Hilty J, Morris RE (2006) Human macrophages do not require phagosome acidification to mediate fungistatic/fungicidal activity against *Histoplasma capsulatum*. J Immunol 176: 1806–13

Nicola AM, Robertson EJ, Albuquerque P, Derengowski Lda S, Casadevall A (2011) Nonlytic exocytosis of *Cryptococcus neoformans* from macrophages occurs *in vivo* and is influenced by phagosomal pH. MBio 2

Otto GP, Nichols BJ (2011) The roles of flotillin microdomains--endocytosis and beyond. J Cell Sci 124: 3933–40

Pike LJ (2006) Rafts defined: a report on the Keystone Symposium on lipid rafts and cell function. J Lipid Res 47: 1597–8

Pike LJ (2009) The challenge of lipid rafts. J Lipid Res 50 Suppl: S323–8

Sainz J, Perez E, Gomez-Lopera S, Jurado M (2008) IL1 gene cluster polymorphisms and its haplotypes may predict the risk to develop invasive pulmonary aspergillosis and modulate C-reactive protein level. J Clin Immunol 28: 473–85

Schmidt H, Vlaic S, Kruger T, Schmidt F, Balkenhol J, Dandekar T, Guthke R, Kniemeyer O, Heinekamp T, Brakhage AA (2018) Proteomics of *Aspergillus fumigatus* conidia-containing phagolysosomes identifies processes governing immune evasion. Mol Cell Proteomics 17: 1084–1096

Schneider A, Rajendran L, Honsho M, Gralle M, Donnert G, Wouters F, Hell SW, Simons M (2008) Flotillin-dependent clustering of the amyloid precursor protein regulates its endocytosis and amyloidogenic processing in neurons. J Neurosci 28: 2874–82

Seider K, Brunke S, Schild L, Jablonowski N, Wilson D, Majer O, Barz D, Haas A, Kuchler K, Schaller M, Hube B (2011) The facultative intracellular pathogen *Candida glabrata* subverts macrophage cytokine production and phagolysosome maturation. J Immunol 187: 3072–86

Simons K, Gerl MJ (2010) Revitalizing membrane rafts: new tools and insights. Nat Rev Mol Cell Biol 11: 688–99

Simons K, Ikonen E (1997) Functional rafts in cell membranes. Nature 387: 569–72

Simons K, Toomre D (2000) Lipid rafts and signal transduction. Nat Rev Mol Cell Biol 1: 31–9

Smith LM, Dixon EF, May RC (2015) The fungal pathogen *Cryptococcus neoformans* manipulates macrophage phagosome maturation. Cell Microbiol 17: 702–13

Solis GP, Hoegg M, Munderloh C, Schrock Y, Malaga-Trillo E, Rivera-Milla E, Stuermer CA (2007) Reggie/flotillin proteins are organized into stable tetramers in membrane microdomains. Biochem J 403: 313–22

Stappers MHT, Clark AE, Aimanianda V, Bidula S, Reid DM, Asamaphan P, Hardison SE, Dambuza IM, Valsecchi I, Kerscher B, Plato A, Wallace CA, Yuecel R, Hebecker B, da Gloria Teixeira Sousa M, Cunha C, Liu Y, Feizi T, Brakhage AA, Kwon-Chung KJ et al. (2018) Recognition of DHN-melanin by a C-type lectin receptor is required for immunity to *Aspergillus*. Nature 555: 382–386

Steele C, Rapaka RR, Metz A, Pop SM, Williams DL, Gordon S, Kolls JK, Brown GD (2005) The beta-glucan receptor dectin-1 recognizes specific morphologies of *Aspergillus fumigatus*. PLoS Pathog 1: e42

Strasser JE, Newman SL, Ciraolo GM, Morris RE, Howell ML, Dean GE (1999) Regulation of the macrophage vacuolar ATPase and phagosome-lysosome fusion by *Histoplasma capsulatum*. J Immunol 162: 6148–54

Stuermer CA (2011) Microdomain-forming proteins and the role of the reggies/flotillins during axon regeneration in zebrafish. Biochim Biophys Acta 1812: 415–22

Thywissen A, Heinekamp T, Dahse HM, Schmaler-Ripcke J, Nietzsche S, Zipfel PF, Brakhage AA (2011) Conidial dihydroxynaphthalene melanin of the human pathogenic fungus *Aspergillus fumigatus* interferes with the host endocytosis pathway. Front Microbiol 2: 96

Triantafilou M, Miyake K, Golenbock DT, Triantafilou K (2002) Mediators of innate immune recognition of bacteria concentrate in lipid rafts and facilitate lipopolysaccharide-induced cell activation. J Cell Sci 115: 2603–11

Tsai HF, Chang YC, Washburn RG, Wheeler MH, Kwon-Chung KJ (1998) The developmentally regulated alb1 gene of *Aspergillus fumigatus:* its role in modulation of conidial morphology and virulence. J Bacteriol 180: 3031–8

Tucker SC, Casadevall A (2002) Replication of *Cryptococcus neoformans* in macrophages is accompanied by phagosomal permeabilization and accumulation of vesicles containing polysaccharide in the cytoplasm. Proc Natl Acad Sci U S A 99: 3165–70

Varshney P, Yadav V, Saini N (2016) Lipid rafts in immune signalling: current progress and future perspective. Immunology 149: 13–24

Vieira FS, Correa G, Einicker-Lamas M, Coutinho-Silva R (2010) Host-cell lipid rafts: a safe door for microorganisms? Biol Cell 102: 391–407

Villalobo A (2018) The multifunctional role of phospho-calmodulin in pathophysiological processes. Biochem J 475: 4011–4023

Volling K, Thywissen A, Brakhage AA, Saluz HP (2011) Phagocytosis of melanized *Aspergillus* conidia by macrophages exerts cytoprotective effects by sustained PI3K/Akt signalling. Cell Microbiol 13: 1130–48

Wojtowicz A, Gresnigt MS, Lecompte T, Bibert S, Manuel O, Joosten LA, Rueger S, Berger C, Boggian K, Cusini A, Garzoni C, Hirsch HH, Weisser M, Mueller NJ, Meylan PR, Steiger J, Kutalik Z, Pascual M, van Delden C, van de Veerdonk FL et al. (2015) IL1B and DEFB1 polymorphisms increase susceptibility to invasive mold infection after solid-organ transplantation. J Infect Dis 211: 1646–57

Woods JP (2003) Knocking on the right door and making a comfortable home: *Histoplasma capsulatum* intracellular pathogenesis. Curr Opin Microbiol 6: 327–31

Zaragoza O, Chrisman CJ, Castelli MV, Frases S, Cuenca-Estrella M, Rodriguez-Tudela JL, Casadevall A (2008) Capsule enlargement in *Cryptococcus neoformans* confers resistance to oxidative stress suggesting a mechanism for intracellular survival. Cell Microbiol 10: 2043–57

Zhang X, Goncalves R, Mosser DM (2001) The isolation and characterization of murine macrophages. In Current Protocols in Immunology, John Wiley & Sons, Inc.

